# Alpha suppression during prehension indicates neural motor drive inhibition

**DOI:** 10.1101/2022.12.02.518923

**Authors:** Oscar Ortiz, Usha Kuruganti, Victoria Chester, Adam Wilson, Daniel Blustein

## Abstract

Changes in alpha band activity (8-12 Hz) have been shown to indicate the inhibition of engagement of brain regions during cognitive tasks, reflecting real-time cognitive load. Despite this, its feasibility to be used in a more dynamic environment with ongoing motor corrections has not been studied. This research used electroencephalography (EEG) to explore how different brain regions are engaged during a simple grasp and lift task where unexpected changes to the object’s properties are introduced. To our knowledge, this is the first study to show alpha activity changes related to motor error correction occur only in motor-related areas (i.e. central areas), but not in error processing areas (ie. fronto-parietal network). This suggests that oscillations over motor areas could reflect inhibition of motor drive related to motor error correction, thus being a potential cortical electrophysiological biomarker for the process, and not solely as a proxy for cognitive demands. This observation is particularly relevant in scenarios where these signals are used to evaluate high cognitive demands co-occurring with high levels of motor errors and corrections, such as prosthesis use. The establishment of electrophysiological biomarkers of mental resource allocation during movement and cognition can help identify indicators of mental workload and motor drive, which may be useful for improving brain-machine interfaces.

**New and Noteworthy:** This research expands on previous fMRI literature by demonstrating that alpha band suppression, an EEG metric with high temporal resolution, occurs over the primary sensorimotor area during error correction of hand movements. This furthers our understanding of alpha suppression beyond processes related to cognitive demands by highlighting how motor control also influences this frequency band. Recognizing that alpha band activity is modulated by both motor and cognitive processes is important in situations where high cognitive demands can lead to a high level of movement errors. Interpretations of such modulation are often attributed only to cognitive demands, whereas a motor process may also play a factor. Furthermore, alpha suppression could be used as a biomarker for error correction with applications in human machine interfaces, such as neuroprostheses.

## Introduction

The human ability to perform fine and precise movement when interacting with objects is an evolutionary development, dating back over 2.5 million years to the first recorded use of tools (Semaw et al., 1997). Disruption to any part of the motor control system can severely affect our lives. For example, stroke patients can suffer hemiparesis and spasticity as a result of damage to motor areas in the brain (Johnson & Westlake, 2021; Yeh et al., 2014). Damage to the peripheral nervous system and end effectors, as in the case of people living with upper-limb loss, can loead to an overall reduction in functionality, resulting in a general reduction in quality of life (McKinley et al., 2007). During interactions with objects, internal models based on the expected properties of the object are used to create predictions about the motor command required and the sensory consequences that arise from the movement (Augurelle et al., 2003; Elias et al., 2008). Errors between predicted and actual sensory consequence can trigger corrective responses that update the motor command to achieve the intended movement and update the sensorimotor system in future interactions (Johansson & Westling, 1988; Shadmehr et al., 2010; Taylor & Ivry, 2011).

Functional magnetic resonance imaging (fMRI) studies have shown that the error between the predicted and actual sensory consequences when lifting an object results in an increased activation of the right inferior parietal cortex, motor cortex, and cerebellum (Jenmalm et al., 2006; Schmitz et al., 2005). Although fMRI evidence sheds light on the brain regions involved in this sensory corrective process, limited time resolution and technological requirements limits its applicability outside of controlled laboratory settings. The fronto-parietal network has also shown differences in alpha activity reflective of the type of grip used to interact with an object (Iturrate et al., 2018). Furthermore, it has also been shown that when the complexity of the sensorimotor action is increased (e.g. more steps), an enhanced suppression of oscillatory power in the alpha band (8-12 Hz) over dorsomedial fronto-parietal areas (i.e. sensorimotor areas) occurs before and during movement execution (Verhagen et al., 2013). Overall, the evidence suggests that activity in the alpha range in the fronto-parietal and motor regions may encode information related to predictive and on-line motor adaptation to the environment.

During cognitive tasks, alpha activity has been demonstrated to be reflective of inhibitory control over specific regions of the cortex, therefore, offering a way to investigate shunting of mental resources from areas that are inhibited (more alpha activity) towards areas that are more task related (less alpha activity), a theory coined the “Gating-by-inhibition” hypothesis (Jensen & Mazaheri, 2010). Recently, a group of researchers (Parr et al., 2019) explored cognitive burden during prosthesis manipulation using measurements of electroencephalography (EEG), specifically focusing on activity of the alpha-band (8-12 Hz) as a proxy for efficient allocation of brain resources. However, the interpretations regarding alpha modulation during this motor task were attributed to cognitive demands, without consideration of motor-control related modulation of this frequency band. Disentangling the interacting effects of motor and cognitive tasks on alpha activity is necessary before the frequency band can be used as a marker of cognitive demand in dynamic settings. Therefore, the purpose of this study was to assess motor-related changes in alpha activity during a reach, grasp and lift movement where unpredictable changes in a custom-made object’s properties were introduced. Using an existing dataset (Luciw et al., 2014), we showed here that alpha activity over the primary motor area is modulated based on corrections of erroneously programmed movements, but activity in the same frequency band over error-corrective areas (fronto-parietal network) did not show a similar modulation. By describing how alpha activity is affected during a simple motor task, we can better understand how cognitive demands modulate movement outside of the laboratory, such as during use of prostheses or brain-machine interfaces.

## Methods

### Experimental Dataset

The study was performed on an open-source dataset (Luciw et al., 2014) that was collected on twelve participants (8 females and 4 males, age range: 19-35 years) performing a precision grasp-and-lift (GAL) of an custom object (Figure 1). The methods used are briefly described below.

**Figure 1:**
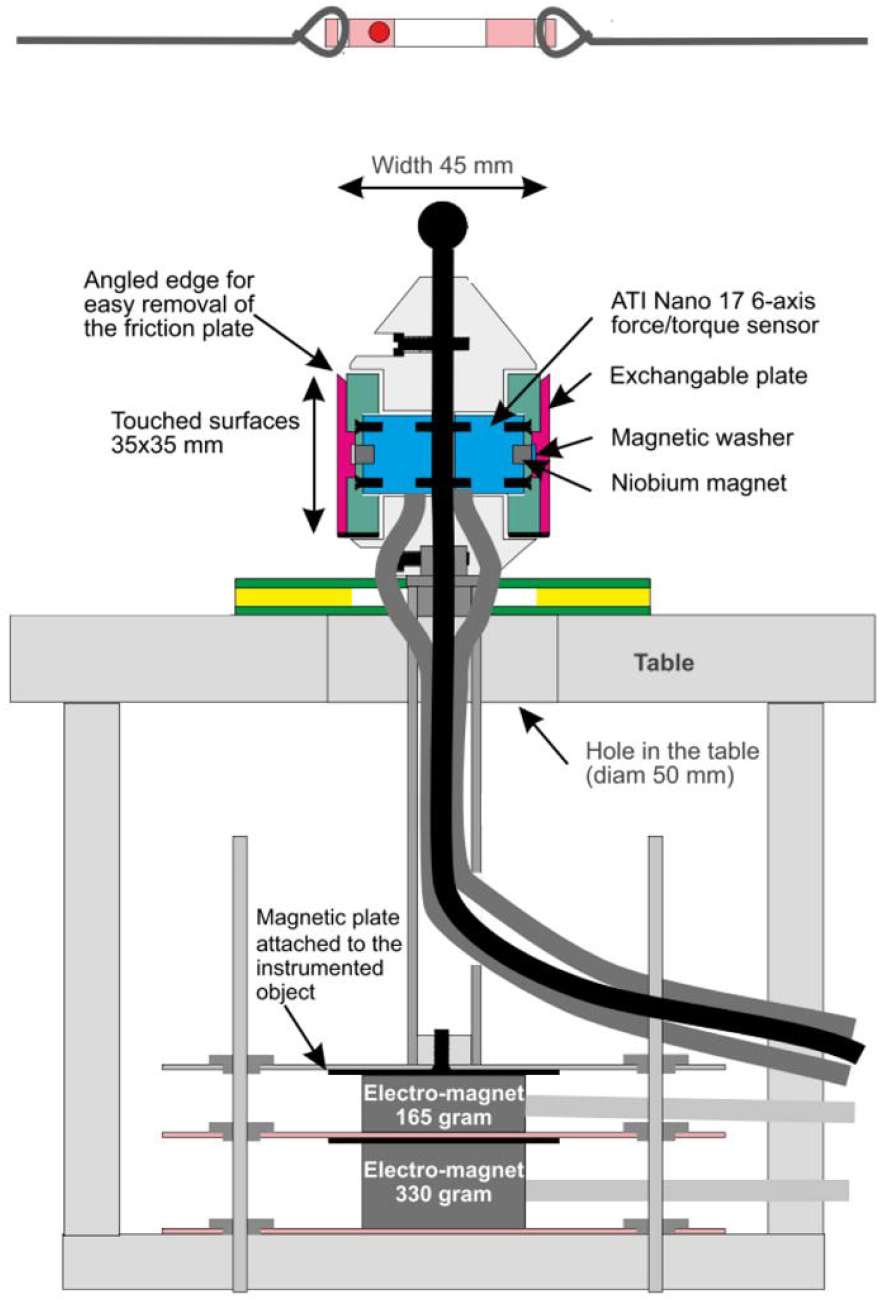
Diagram of object used in Reach and Grasp trials (from Luciw, Jerocka & Edin, 2014).

### Data Acquisition and Instrumentation

During the experiment, EEG, EMG, position and force data were recorded. A 32-electrode EEG system using the standard 10-20 positions was used to record brain activity at 500 Hz (ActiCap, Brain Products, Gilching, Germany). Five EMG sensors sampled muscle activity at 4 kHz from arm muscles including the anterior deltoid (AD), brachioradialis (BR), flexor digitorum (FD), common extensor digitorum (CED), and the first dorsal interosseus (FDI) muscles (Figure 2). EMG was recorded using preamplifiers (bandwidth 6Hz-2 kHz) mounted on the skin directly above the muscle. Electrodes were 2mm in diameter and 12 mm apart. Four 3D position sensors (FASTRAK, Polhemus Inc, USA) recorded the position (XYZ Cartesian coordinates) and orientation (azimuth, elevation, and roll) of the object, index finger, thumb, and the wrist at 500 Hz. Finally, the surface plates of the object were coupled to force transducers that recorded 3D forces and torques at 500 Hz.

**Figure 2:**
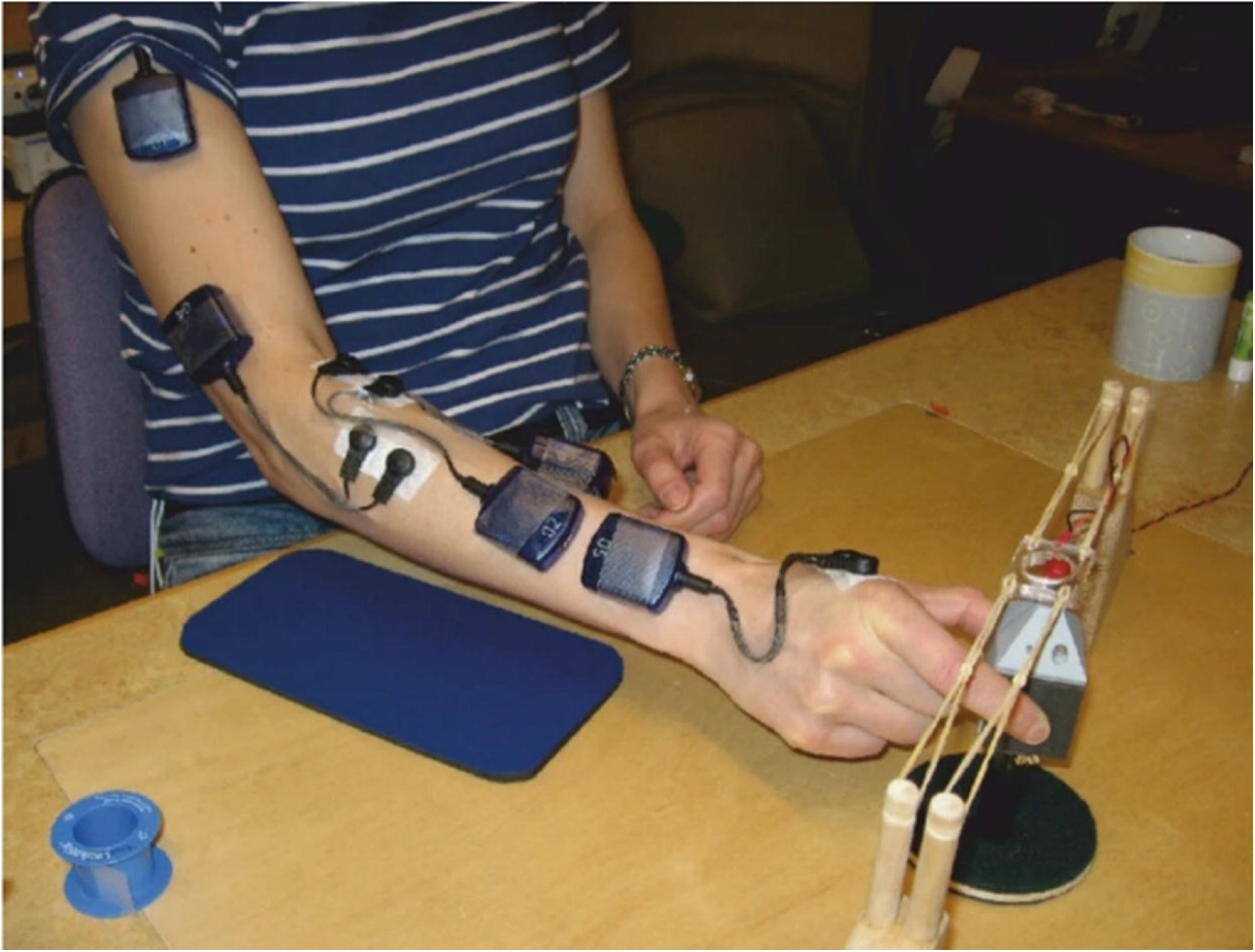
Experimental set-up for the Reach and Grasp Experiment (from Luciw, Jerocka & Edin, 2014).

### Task Description

Participants were asked to start the movement when an LED on a graspable object (width between plates: 45 mm, surface area: 35X35 mm) flashed red (Figure 1). Participants were instructed to begin each movement with their right wrist resting on the table so that the forearm was suspended over the edge of the table. They were instructed to begin their movement by first reaching from the initial position towards the object, grasp the object with the thumb and index finger, and lift it so that the top of the object was within a circle target suspended about 5 cm above the object. Participants then were instructed to place the object back down when the LED turned off (∼2 seconds after the LED turned on) and return their arm to the initial position in preparation for the next lift.

Each participant performed two different experimental series lifting a custom object whose weight and surface friction could be altered without the participant’s knowledge (Figure 1, for a more detailed description, see Luciw et al., 2014). The weight series involved 34 lifts with 12 unpredictable weight changes (between 165, 330, and 660 g). There were six different weight series schedules, so each weight was repeated 1-4 times (expected) and then suddenly changed (unexpected). The friction or surface series involved 34 lifts with variable surface friction (sandpaper, suede, or silk) and similarly, the same texture was presented 1 to 4 times and then changed unexpectedly. All sequences and changes were balanced across the constructed series. Each participant performed 6 weight series (each series consisted of 34 lifts with 12 unpredictable changes to the weight) and 2 friction series (34 lifts including 12 unexpected changes to the surface friction) amounting to 3,264 lifts that were included across all participants in the analysis. See Luciw et al, 2014 for details about the object and paradigm.

### Data Analysis

#### EEG

Signals were average-referenced and band-passed filtered from 1 to 45 Hz using a zero-phase (two-pass) FIR filter of order 500. Independent Component Analysis and visual inspection of the signals was performed to remove components accounting for blinks, eye movements and other non-neural activity (Delorme & Makeig, 2004). The continuous data were then epoched and time locked from 500 ms before and 1000 ms after initial contact with the object. Initial contact was defined as the moment when the summed forces perpendicular to the grip platforms on the object reached two standard deviations from the baseline period. Changes in alpha activity were computed by calculating the mean change in spectral power (in dB) from baseline (-500 to 0 ms) for different frequencies and latencies using a complex Morlet wavelet transform (Tallon-Baudry et al., 1997; Herrmann et al., 1999). The number of cycles was selected according to the frequency and was increased from 0.5 to 13.8 for a frequency range of 1-30 Hz. The baseline power spectrum was calculated for the 500 ms baseline period. Changes in the alpha (8–12 Hz) frequency band in dB in for each subject were calculated for the analysis, and power values were averaged over trials to derive the power spectral estimation. After the time-frequency decomposition, alpha activity (8-12 Hz) was averaged across several regions of interest: left temporal (T7), left central (C3), frontal (F3, Fz, F4), right central (C4), right temporal (T8), parietal (P3, Pz, P4), and occipital (O1, O2).

Data were grouped differently for two different comparisons. To test the effect of expectation alone over error processing regions, trials were sorted into two categories, trials where the properties were ‘expected’ (i.e. same weight or same friction condition than the previous one) and trials where properties were ‘not expected’ (i.e. different weight or friction condition as previous one). To test whether alpha activity over the primary motor area follows changes in motor drive based on on-line corrections to erroneously programed lifts, trials were grouped into two conditions – trials programed for lower weight than required (i.e. a heavier trial after a series of lighter trials) and trials programmed for a heavier weight than required (i.e. a lighter trial after a series of heavier trials).

#### Position, Force Data and EMG

Position of the object as well as lift and grip force data were used to evaluate motor behavior during the task. Signals were low-pass filtered at 9 Hz using a zero-phase (two-pass) FIR filter of order 500. Derivatives of position and force were also extracted to investigate rate of force development and velocity. To quantify the differences between conditions, grip-force rate (GFR), and lift-force rate (LFR) were extracted from the filtered signals coming from the force transducers of the object at two time points: at the moment of lift-off (defined as the moment where the horizontal component of the object’s position increased 2 standard deviations from baseline) and at the moment where they reached their maximum value during the initial second of the lift.

For the EMG recordings, 1 second of data was epoched from the beginning of the loading phase for analysis for each of the muscles being recorded. A second order Butterworth bandpass filter from 20 to 500 Hz was applied to the signals. The signal was then rectified, and the envelope of that signal was computed as the magnitude of its analytic signal. To compare changes before and after initial contact with the object, the data were then normalized by subtracting the mean baseline value of the EMG between -200ms and -500ms prior to initial contact with the object. As with position and force, quantification of muscle activity was performed by extracting the peak EMG value from the enveloped signal at the moment of lift-off as well as the maximal EMG value within the first second of the lift. A summary of all the outcomes measured can be found in Table 1.

**Table 1:** Summary of outcome measures for each measurement device.

| System | Outcome measures being recorded |
| --- | --- |
| EEG | Alpha activity across the seven Rols in windows of 100 ms starting at initial object touch. |
| EMG | Envelope activity from the five muscles being recorded during lift-off and at the maximal activation within the initial second of the lift. |
| Kinematic and Kinetic Traces | Rate of grip force, rate of lift force, and vertical velocity of the object during lift-off and at the maximal value of the trace within the initial second of the lift. |

## Statistical Analysis

### EEG

#### The effect of expectation on alpha activity

A repeated-measures ANOVA was conducted over the error processing regions using a 2 x 2 x 10 factor design with expectation (expected, unexpected), region of interest (RoI) (parietal, frontal) and time window (baseline-100 ms in steps of 100 ms) for the weight and friction data, separately.

#### The effect of object property on alpha activity

A repeated-measures ANOVA was conducted over motor drive electrodes using a 2 x 2 x 10 factor design with property expectation (heavier than expected, lighter that expected for weight; and less surface friction than expected or more surface friction than expected for friction series), RoI (left central, right central), and time window (baseline-100 ms in steps of 100 ms) for the weight and friction data, separately. Degrees of freedom were adjusted with the Greenhouse-Geisser correction if the sphericity assumption was violated as indicated by a Mauchly test. Pairwise t-tests with Bonferroni corrections were used for post-hoc analysis. In all analyses, a significant criterion of α= 0.05 was used.

### Position and Force Data and EMG

#### Effect of expectation on different conditions

Pairwise comparisons with Bonferroni corrections were performed among the metrics of motor behavior: lift-off and maximal GFR and LFR values, and EMG envelope values (see data analysis section for details). To isolate the effect of expectations and avoid confounding variable weights or surface frictions, contrasts were performed between objects lifts that were performed with the same weight or surface friction, but only varied in weight and surface friction in the previous lift. For example, to determine the difference between expected weight or higher than expected weight, lifts with the same weight were compared (i.e. 660 g), but the expected was preceded with a lift of the same weight (i.e. 660 g → 660 g) and the unexpected heavy condition was preceded with a lift of lighter weight (i.e. 165 g → 660 g).

## Results

### EEG

#### The effect of expectation on alpha activity in error processing regions

A repeated-measures ANOVA revealed a main effect of time point (*F*(10, 110) = 3.06, *p* < 0.05) and an interaction effect between RoI and time point (*F*(10, 11) = 3.48, *p* < 0.001) for the weight series only. However, no main effect of expectation (i.e. trials with expected weight vs. unexpected weight) was found for the weight (*F*(1, 11) = 0.39, *p* = 0.54), or the friction (*F*(1, 11) = 0.71. *p* = 0.41). These results suggest that the alpha activity over error processing regions was not able to reflect error correction during a motor task.

#### The effect of object property on alpha activity in motor regions

The analysis revealed a main effect of time point for the weight (*F*(10, 110) = 4.34, *p* < 0.05) and friction series (*F*(10, 110) = 2.20, *p* < 0.05), respectively. Furthermore, the friction series showed an interaction effect of RoI and time point (*F*(10, 110) = 2.94, *p* < 0.05). The weight series showed an interaction effect between the property expectation and time (*F*(20, 220) = 2.70, *p* < 0.001) and a three-way interaction of expected grip force required, RoI, and time (*F*(20,220) = 1.78, *p* < 0.05). Post-hoc analysis showed that the left central region responsible for the control of the contralateral movement showed a greater alpha activity in trials programmed for heavier weights than expected (600 g to 165 g) compared to trials programmed for a lower weight than expected (165 g to 600 g) between 700 ms (*t*(11) = -3.35, *p* < 0.05) and 800 ms (*t*(11) = -2.48, *p* < 0.05) (Figure 3).

**Figure 3:**
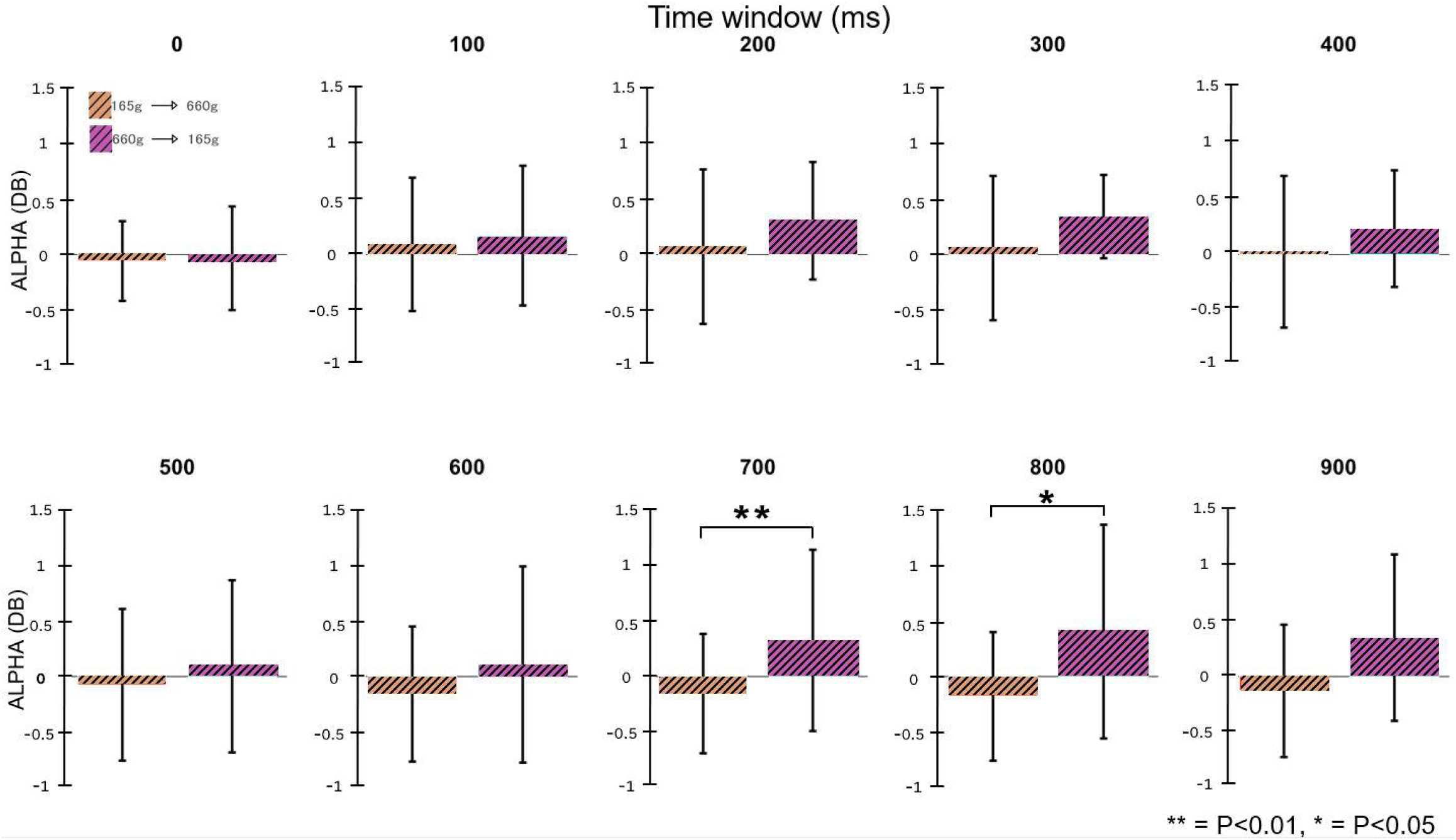
Time analysis during weight series trials over the left central region for lifts with an unexpected weight drop (660 to 165 g, purple) and those with an unexpected weight increase (165 g to 660 g, brown). Activity in the 700 to 800 ms windows were significantly greater in the trials with lower weights than expected.

### EMG

Mean traces of the EMG envelope are shown in Figure 4 and 5 for the weight and friction series, respectively. *<u>Weight series:</u>* Compared to trials where force was properly gauged, lifts performed with a weight lighter than expected showed a significantly greater EMG activation over the BR (*t*(11) = -2.71, p=0.020), and CED (*t*(12) = -4.15, p=0.0016) during lift off, as well as a significantly greater maximal activity within one second after object touch over the AD (*<u>t</u>*(11) = -2.81, p=0.017), BR (*t*(11) = -2.20, p=0.049), CED (*t*(11) = -2.78, p=0.018), FD (*t*(11) = -2.78, p=0.018), and FDI (*t*(11) = - 2.35, p=0.038) muscles (Figure 6. A, C, E, G, I, purple bars). Furthermore, during lifts with a weight higher than expected, participants showed a significantly lower activity of the AD (*t*(11) = -3.33, p=0.0067) during lift-off, and as well as a significantly greater maximal activity within one second of initial object touch in the BR (*t*(11) = 4.28, p=0.0013), FD (*t*(12) = 2.51, p=0.029), and FDI (*t*(11) = 2.81, p=0.0170) muscles (Figure 6. A, C, E, G, I, brown bars). *<u>Friction series:</u>* No significant changes in EMG activity were found across the muscles studied (Figure 6. B, D, F, H, J).

**Figure 4:**
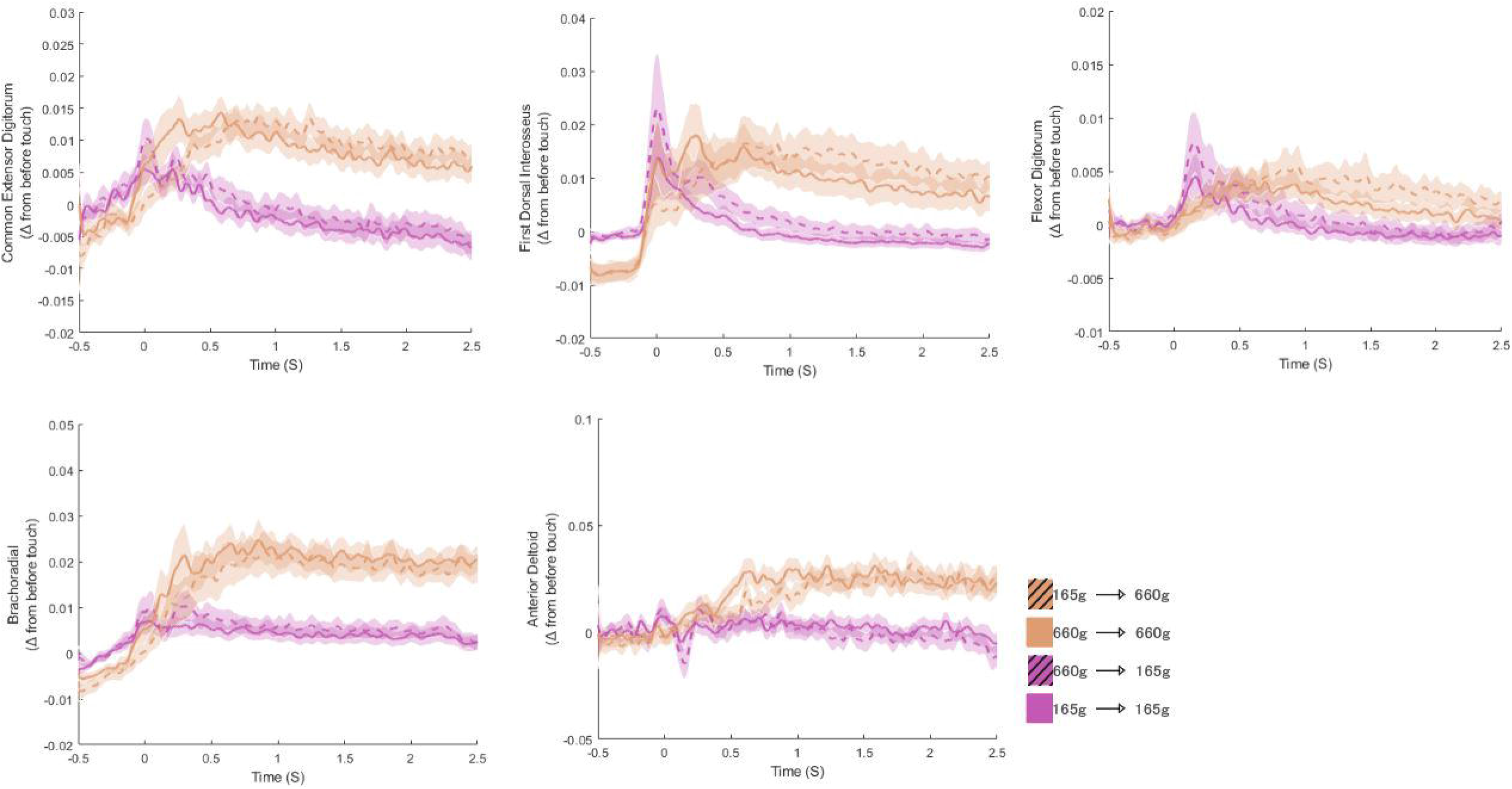
EMG traces for the Common Extensor Digitorum, First Dorsal Interosseous, Flexor Digitorum, Brachioradialis, and Anterior deltoid during the weight series trials. Shaded area shows the 95% confidence interval. Data are divided in colors by weight being lifted (brown = 660g, purple = 165g) and expectation (expected = solid, unexpected = hashed).

**Figure 5:**
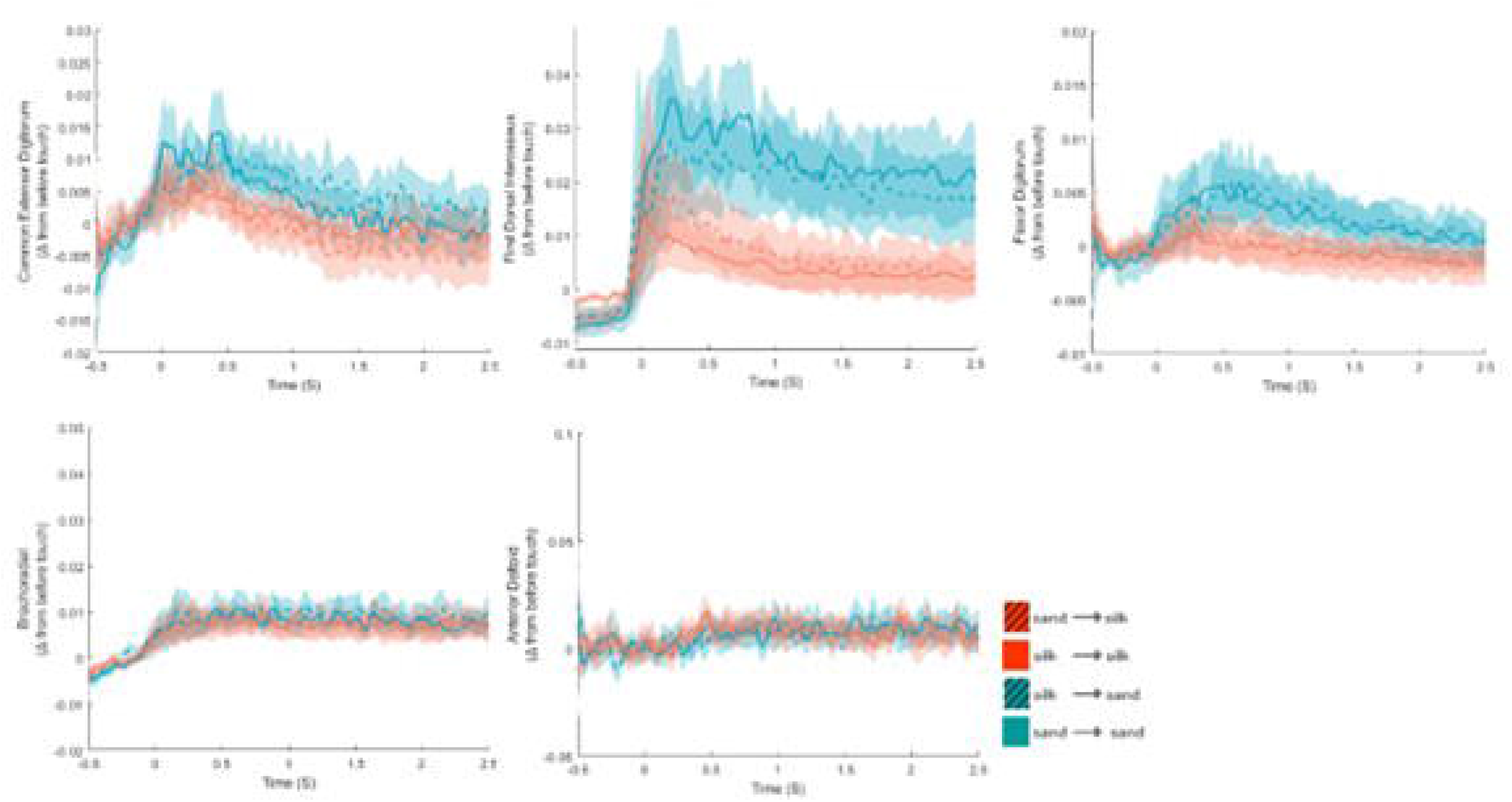
EMG traces for the Common Extensor Digitorum, First Dorsal Interosseous, Flexor Digitorum, Brachioradialis, and Anterior deltoid during the friction series trials. Shaded area shows the 95% confidence interval. Data are divided by the lifted object’s surface texture (silk = red, sandpaper = blue) and texture expectation (expected = solid, unexpected = hashed).

**Figure 6:**
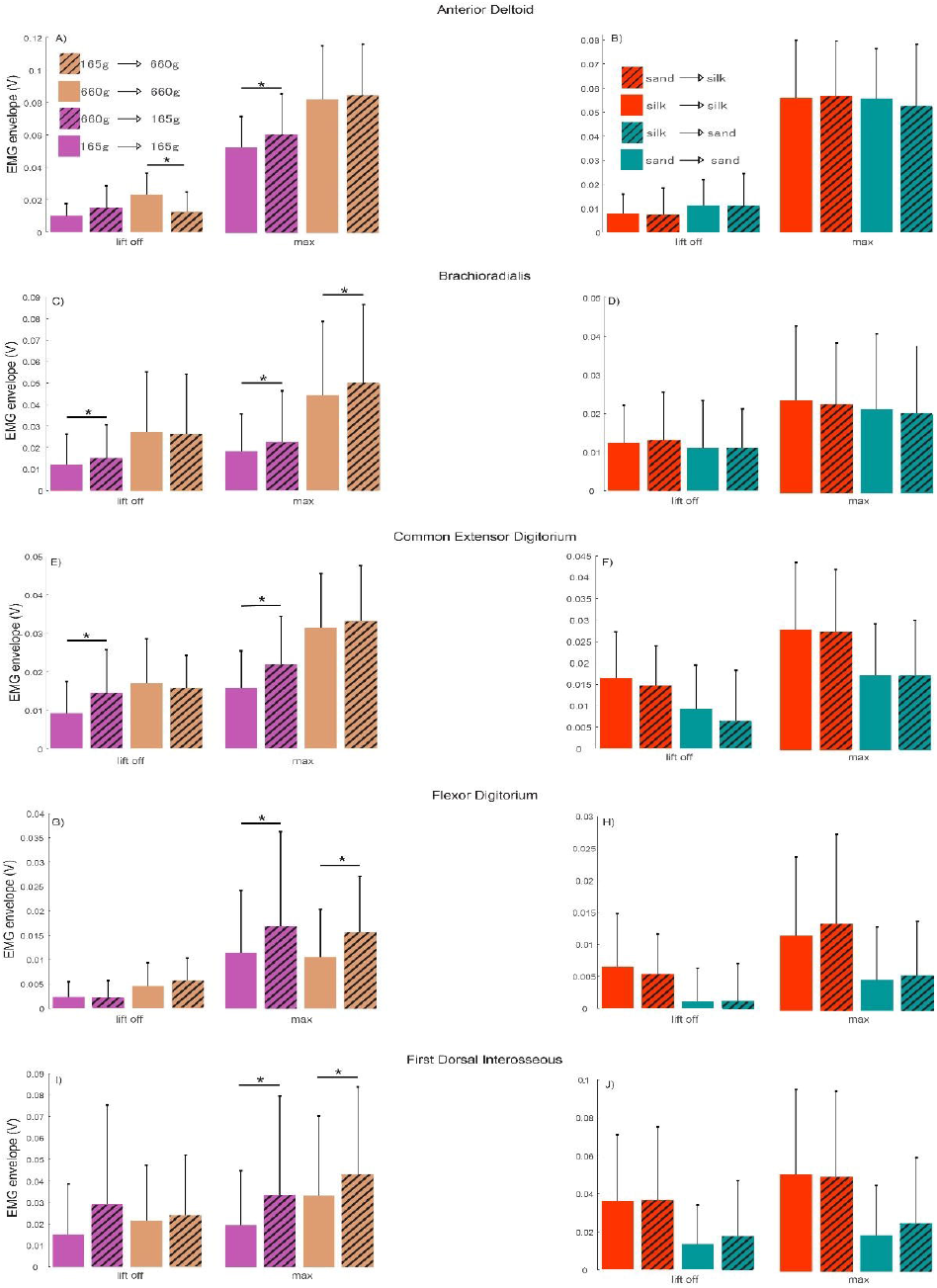
Summary of results for all EMG muscles during lift-off and maximal level of activity for the weight (left column) and friction (right column) series trials. Data from the weight series (left) are divided in colors by weight being lifted (brown = 660g, purple = 165g) and expectation (expected = solid, unexpected = hashed). Friction data (right) are divided by surface texture (silk = red, sandpaper = blue) and expectation (expected = solid, unexpected = hashed).

### Position and Force Data

Mean traces for the position and force data are shown in Figures 7 and 8 for the weight and friction series, respectively. *<u>Weight series:</u>* Kinetic analysis showed significantly greater GFR at lift-off (*t*(11) = -6.35, *p* < 0.05) and maximal points (*t*(11) = -7.17, *p* < 0.05) Figure 6. A) and significantly greater LFR (*t*(11) = -2.54, *p* < 0.05) at lift-off (Figure 9. C) when the lift was programed for a greater weight than expected (600 g to 165 g) compared to when it was expected (165 g to 165 g). During lifts with an unexpected weight increase, we found significantly lower GFR at lift-off (*t*(11) =2.23, *p* < 0.05) and maximal point (*t*(11) = 6.98, *p* < 0.05) (Figure 6. A) and a significantly lower maximum LFR (*t*(11) = 5.21, *p* < 0.05) (Figure 9. C) compared to when there was no weight change (660 g to 660 g).

**Figure 7:**
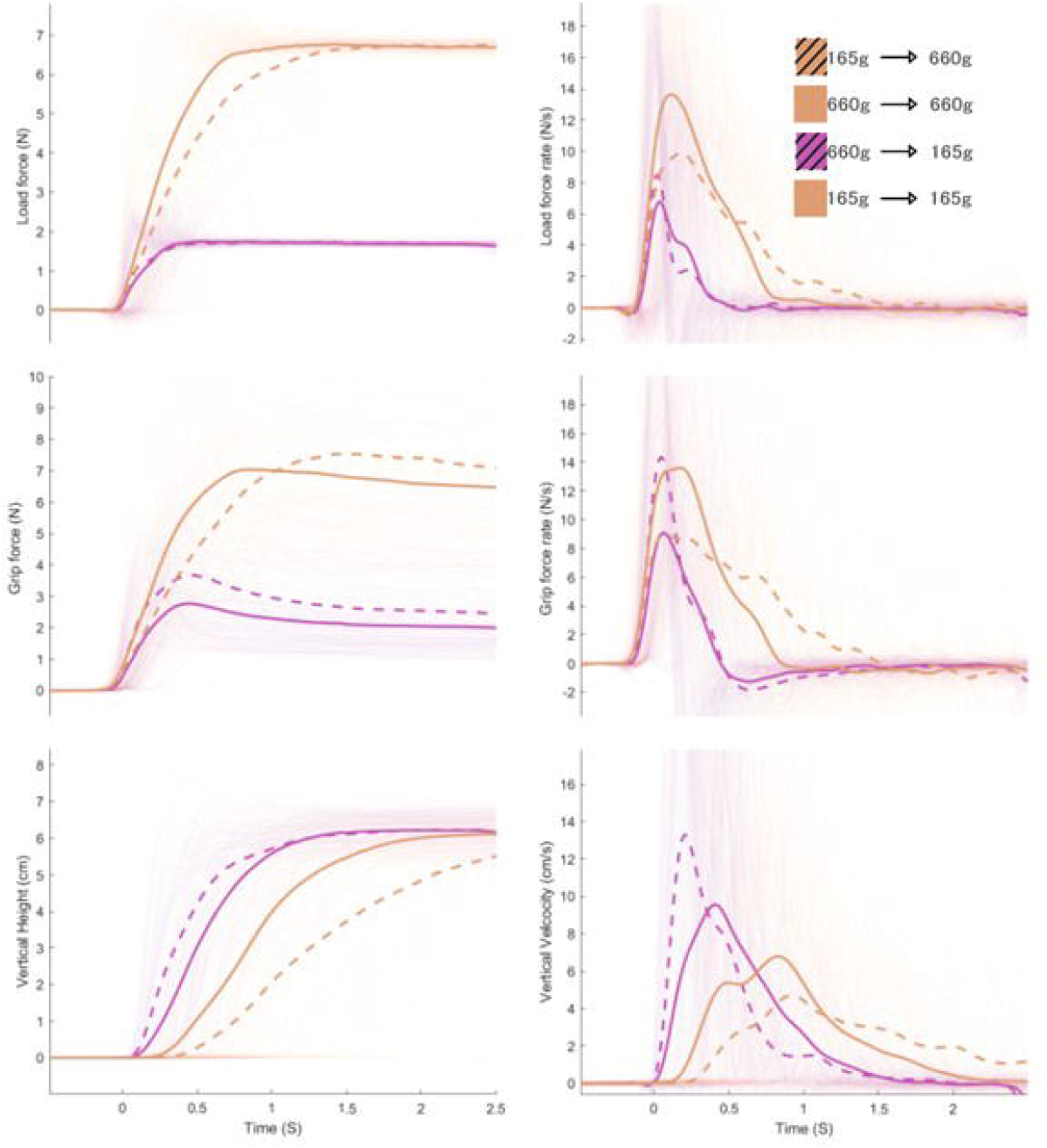
Kinematic and kinetic traces of the weight series. Panels on the left represent the load force (top row), grip force (middle row) and vertical height (bottom row), with their respective derivatives presented in the right column, for each of the experimental conditions.

**Figure 8:**
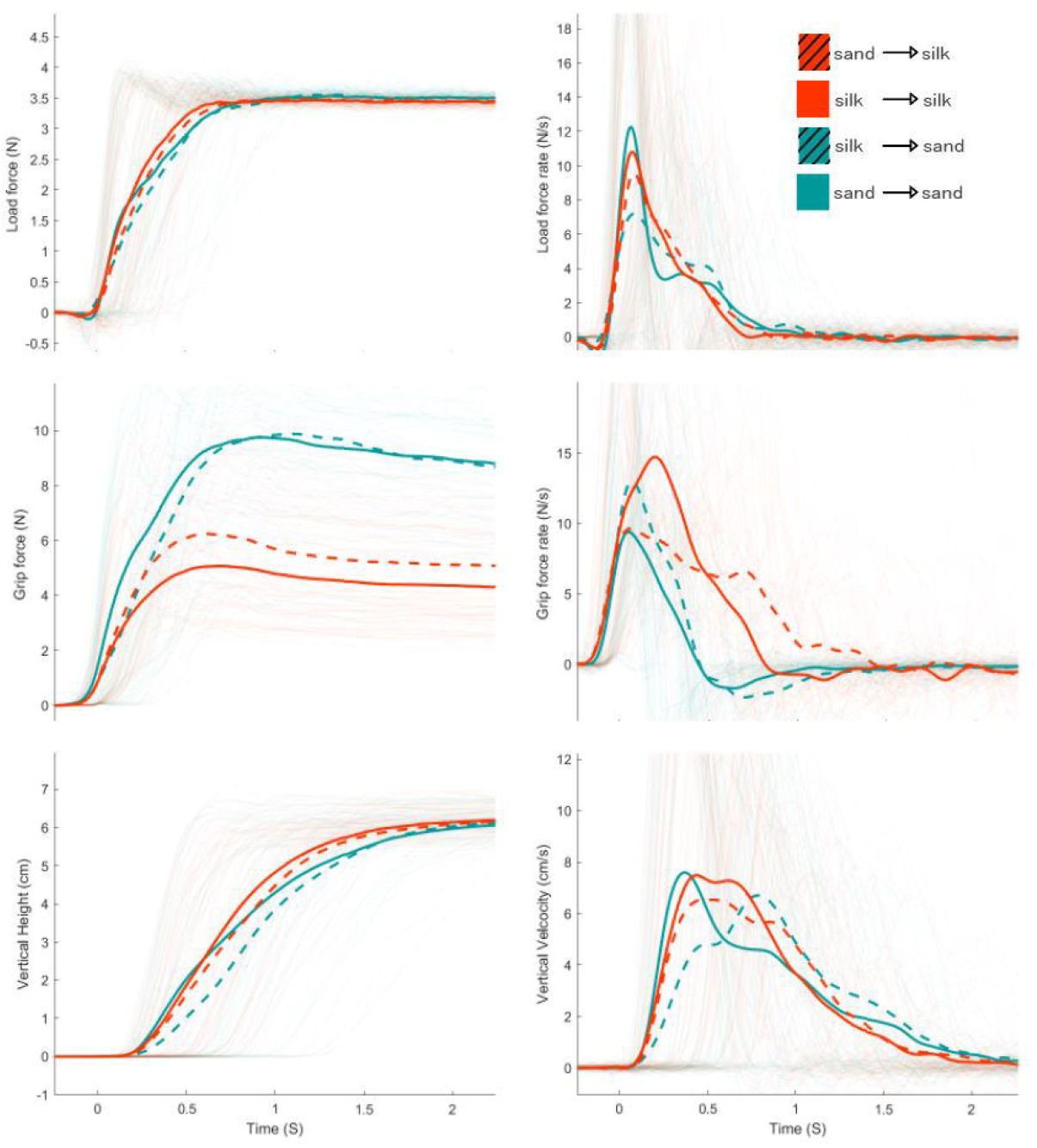
Kinematic and kinetic traces for the friction series. Panels on the left, from top to bottom, represent the load force, grip force and vertical height (and their respective derivatives on the right) for each of the experimental conditions.

**Figure 9:**
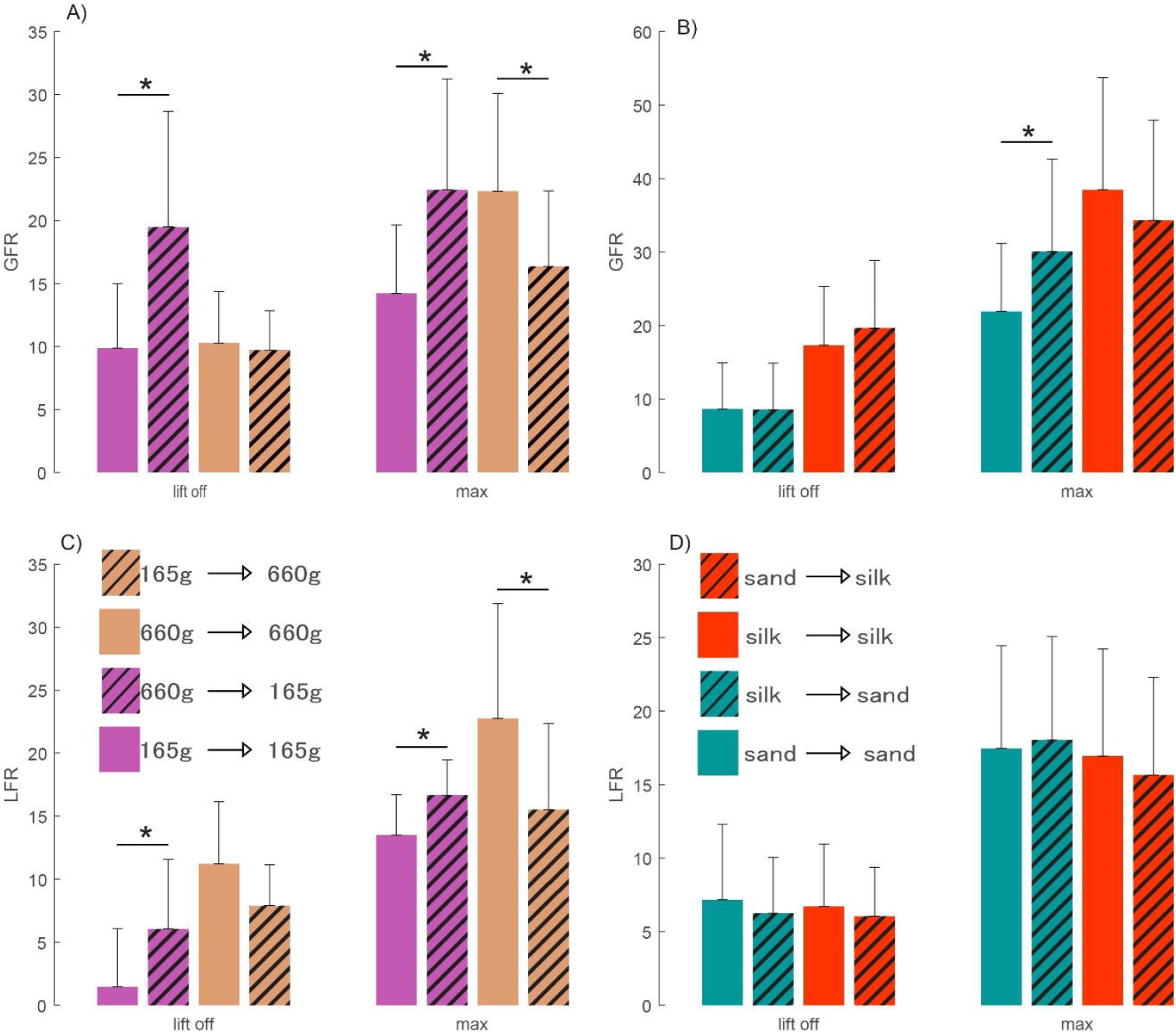
Summary of the effect of expectation on grip force rate (GFR, A and B) and lift force rate (LFR, C and D) during weight and friction series trials. Lift off values were extracted when the object was first lifted from the table and the maximum values were extracted at the peak of each of the rates of force (see Figures 7 and 8). Data from the weight series (left) are divided in colors by weight being lifted (brown = 660g, purple = 165g) and expectation (expected = solid, unexpected = hashed). Friction data (right) are divided by surface texture (silk = red, sandpaper = blue) and expectation (expected = solid, unexpected = hashed).

#### Friction series

Maximal GFRs (Figure 9. B) for trials programmed for silk but performed with sandpaper (silk to sandpaper) were significantly greater (*t*(11) = -2.96, *p* < 0.05) than those when sandpaper was expected (sandpaper to sandpaper). No other significant differences were found in behavioral analysis of the friction series.

## Discussion

We investigated differences in alpha activity over error-detection and motor drive regions of the brain during the grasping and lifting of a modifiable object. Kinematic, kinetic and EMG data show that the expectation of object properties affects the strategy being used to interact with the object, primarily in the weight domain. We see evidence of different muscle activity and force developing patterns depending on the properties of the previous object moved. Light weight trials preceded by heavier trials showed greater FD and FDI activity than when the following trial did not change weight. This change in muscle activation was accompanied by a greater GFR and LFR suggesting that the weight properties of previous lifts influence the mental model used when lifting the object (Saunders & Vijayakumar, 2011). By overestimating the force required to lift the object, the forward model used to estimate the object properties generates a motor command that requires correction to account for the present object properties (Augurelle et al., 2003; Baugh et al., 2012; Johansson & Westling, 1988).

Due to the effect of previous lift outcomes on the internal model of the object, we expected engagement of motor error correction regions of the brain. Contrary to this, however, we did not observe any significant differences over the frontal or parietal areas between the expected and unexpected trials. These areas are known to be part of the error correction system and have been related to updating motor commands based on the discrepancy between actual and predicted sensory information (Blakemore & Sirigu, 2003). Furthermore, they are also related to the updating of sensorimotor memories of an object’s physical properties (Ehrsson et al., 2003; Jenmalm et al., 2006). Other behavioral fMRI studies have also linked activity over this region in tasks involving manipulation, lifting, and pinching of objects (Binkofski et al., 1999).

One possible reason for the lack of observed differences in activity in error correction regions is that due to the ambiguity of the object’s properties and the high number of repetitions, participants might have adapted a feedforward strategy that minimizes the amount of movement errors while lifting. Indeed, previous work suggests that, in the face of sensorimotor uncertainty, this is the preferred strategy over one that selects the most likely prediction of the object’s properties (Cashaback et al., 2017). With this strategy, the nervous system would have to build a representation of the environmental uncertainty to develop a “point estimate” of a single weight based not only on the immediate previous lift, but all lifts preceding the current one (Kording & Wolpert, 2004; Wolpert et al., 1995). In this case, it is possible for that error integration signal to be attenuated since the strategy aims to minimize the error between the estimate and the possibility of all weights or friction surfaces.

Another possible explanation for the lack of differences in alpha activity in the error correction region surrounds the gating-by-inhibition hypothesis assumptions of the frequency content of the inhibition (Jensen & Mazaheri, 2010). This framework suggests that alpha activity represents a pulsed inhibition that can reduce the capabilities of information processing of a given area. Physiologically, the pulsing inhibition has been linked to GABAergic (inhibitory) feedback inputs from interneurons that have been shown to modulate this specific band (Jones et al., 2000; Lörincz et al., 2008). Evidence for this inhibitory activity reflected by alpha oscillations comes from changes in brain activity related to cognitive processes, such as a memory tasks (Bashivan et al., 2014; Rottschy et al., 2012) or visual field attention (Kelly et al., 2009; Rihs et al., 2007). It is possible that the timescale of the inhibition over this area related to motor activity diverges from inhibition related to cognitive tasks, and subsequently, any pulsed inhibition through interneurons may not be reflected by changes to alpha rhythm alone. . These differences in time-scales might be why changes in activity appear in this area in fMRI studies but not when examining EEG alpha activity. This possibility could further be explored through analysis of motor-error related differences in parietal activity over different frequency bands and time scales, but it is outside of the scope of this initial analysis focusing on alpha-related changes.

Another possible explanation of these findings is that the spatial resolution of the EEG setup during the experiment was not specific enough to detect changes related to motor activity. The right supramarginal gyrus identified previously as involved in error correction (Jenmalm et al., 2006) is only one of many gyri found in the parietal lobe (Braver et al., 1997; Ghosh & Gattera, 1995; Neal et al., 1990). EEG systems are prone to poor spatial precision because of volume conduction across the scalp and the low number of electrodes (i.e. 32 electrodes) in standard setups. It is possible that through the averaging of the parietal electrodes, the specific activity over this sub-region of the parietal lobe might have been lost. Future studies could address the spatial resolution issue by using high-density EEG systems containing up to 256 electrodes and performing single electrode analysis as opposed to functional region analysis to increase the spatial localization capabilities of this neuroimaging technique (Barzegaran & Knyazeva, 2017), However, from a practical perspective, using highly variable biomarkers that require high density EEG might not be feasible when applied in real world scenarios due to practical constraints such as set-up time and drift in electrode connectivity over time (Gentili et al., 2014; Jaquess et al., 2018). Overall, our results do not support the role of alpha activity as a biomarker for processes involving the parieto-frontal network, and thus highlighting the limitations of using alpha activity maps to shape the functional architecture of the brain during manual tasks.

When comparing activity over the central regions we did observe significantly higher alpha activity over the left central hemisphere in trials programmed for higher than expected weights compared to trials programmed for lower than expected weights. In these trials we also saw a difference in force production patterns. In comparison to their expected trials, we saw a reduction in peak GFR and LFR in trials programmed for a higher weight than necessary but an increase in GFR and LFR in trials programmed for a lower weight than necessary. Taken together, the results show a relatively higher alpha activity for trials where grip force and lift force were reduced during the lift in comparison to a lower relative alpha activity where grip and lift force were increased during the lift. As alpha is thought to reflect neuronal patterns that reflect inhibition, this could suggest that the higher alpha activity during lifts with unexpected lower weights might reflect the on-line inhibition over motor areas to reduce the neural drive to the muscles and thus reduce force produced.

Our results show a similar pattern of activation as previously reported in an fMRI study where lifts programmed for higher weights than expected demonstrated a lower BOLD signal in the left central region compared to trials programmed for a higher grip force (Jenmalm et al., 2006) which also concluded that these results reflect a reduction of neuronal drive. Despite this, our EMG results show that the finger flexors, FDI and FD, showed a spike during trials that were lighter than expected compared to when the weight was expectedly light which then merged onto the same level of activation later in the lift (Figure 4). This spike in muscle activity required to grip the object could be a compensatory response to avoid slipping of the object due to the higher rates of vertical movement (Figure 8, vertical velocity). However, both FDI and FD EMG traces merge to match the levels of activation when the object’s weight is expected. It is important to note that these spikes happened early during the lift (∼300 ms, Figure 4), whereas alpha suppression appears later during the lift (∼700-800 ms). From the timing of the results, the alpha suppression could be reflective of the inhibition of this corrective EMG spike to return to the correct movement pattern.

The higher alpha activity over the contralateral central region during movements programmed for higher weights might reflect the pulsed inhibition experienced by that region of the brain to adapt the grip force on-line during the lifting portion of the movement. Mechanistically, this would align with the GABAergic (inhibitory) feedback inputs from interneurons that have been shown to modulate this specific band during mental tasks (Jones et al., 2000; Lörincz et al., 2008). This suggests that there might be some overlap in motor-related inhibition of the primary motor cortex with inhibition related to other cognitive processes. Future studies looking to use the gating-by-inhibition framework to study how different cognitive paradigms are reflected in the alpha band should be mindful of the potential overlap of this motor-related inhibition over the primary motor cortex, as activity over this area could be reflective of motor inhibitory mechanisms.

A novel aspect of the present work is the ability to extract the timing of motor drive inhibition in the brain. The onset of the relative enhancement of inhibitory alpha activity over the left central region during lifts programmed for higher weights occurred around 700 to 800 ms after the initial touch. Due to limitations in temporal resolution (∼ 1 s), previous fMRI research reporting this pattern of activation has not been able to estimate the timing of this event, limiting their ability to explicitly state that the increase in brain activity in the left central region reflects an on-line mechanism implemented by the participant to reduce the motor output in a compensatory manner (Jenmalm et al., 2006; Schmitz et al., 2005). In this study, alpha activity over the left central region diverged well after the initial touch of the object. This pattern of activity over motor areas may reflect the on-line reduction of neural drive to the hand musculature for corrective responses.

To our knowledge, this is the first evidence of error-correction related alpha modulation demonstrated on EEG data over motor regions during a manual task, expanding on previous fMRI studies. Furthermore, the data presented here shows that modulation over the alpha band does not occur over error correction regions. These findings suggest that not all engagement of task-relevant regions is reflected through reduced alpha activity, and moreover, explores the notion that the pulsed inhibition mechanism expands beyond cognitive processes and could also reflect corrective motor-drive changes. Thus, if advances in cognitive load research and assessment are to be applied outside of the laboratory, efforts must be made to bridge cognitive and motor studies as they relate to using EEG to shape the brain’s architecture, as motor and cognitive brain activity are likely overlapping. This carries implications about how future research efforts interpret cognitive load results as they pertain to alpha activity, as motor-related inhibition can also contribute to the activity measured, especially during tasks with high rates of error correction such a prosthesis use. In conclusion, this study showed evidence that alpha activity over motor drive regions can be modulated based on error correction during unexpected changes of an object being manipulated.

